# Chromatin accessibility landscape and active transcription factors in primary human invasive lobular and ductal breast carcinomas

**DOI:** 10.1101/2022.04.08.487589

**Authors:** Sanghoon Lee, Hatice Ulku Osmanbeyoglu

## Abstract

**Background:** Invasive lobular breast carcinoma (ILC), the second most prevalent histological subtype of breast cancer, exhibits unique molecular features compared with the more common invasive ductal carcinoma (IDC). While genomic and transcriptomic features of ILC and IDC have been characterized, genome-wide chromatin accessibility pattern differences between ILC and IDC remain largely unexplored.

**Methods:** Here, we characterized tumor-intrinsic chromatin accessibility differences between ILC and IDC using primary tumors from The Cancer Genome Atlas (TCGA) breast cancer assay for transposase-accessible chromatin with sequencing (ATAC-seq) dataset.

**Results:** We identified distinct patterns of genome-wide chromatin accessibility in ILC and IDC. Inferred patient-specific transcription factor (TF) motif activities revealed regulatory differences between and within ILC and IDC tumors. EGR1, RUNX3, TP63, STAT6, SOX family, and TEAD family TFs were higher in ILC, while ATF4, PBX3, SPDEF, PITX family, and FOX family TFs were higher in IDC.

**Conclusions:** This study reveals the distinct epigenomic features of ILC and IDC and the active TFs driving cancer progression that may provide valuable information on patient prognosis.

## Introduction

Breast cancer, the leading malignancy in women, has molecularly discrete subtypes based on the expression of estrogen receptors alpha (ESR1, also known as ER), progesterone receptors (PGR, also known as PR), and/or the amplification of human epidermal growth factor receptor 2 (*ERBB2*, also known as HER2). Of the ~200,000 newly diagnosed cases of invasive breast cancer each year, 70% are estrogen receptor-positive (ER+) (1). More patients die from advanced ER+ breast cancer than all other breast cancer types combined. ER+ breast cancer comprises two main histological subtypes with varying molecular features and clinical behaviors: 85–90% are invasive ductal carcinoma (IDC) and 10–15% are invasive lobular carcinoma (ILC) (2–4). ILC is predominantly ER+ and PGR-positive but can, rarely, show HER2 protein overexpression. While ILC is initially associated with longer disease-free survival and a better response to adjuvant hormonal therapy than IDC, the long-term prognosis for ILC is worse than IDC; 30% of ILC patients will develop late-onset metastatic disease up to 10 years after the initial diagnosis (5). In several retrospective studies that compared clinical and pathological responses, ILC also appeared less responsive to chemotherapy than IDC (6–8).

Although ER+ ILC and IDC tumors are treated clinically as a single disease (9), recent studies have established ER+ ILC as a distinct disease with unique sites of metastasis, frequent presentation of multifocal disease, delayed relapses, and decreased long-term survival compared to ER+ IDC tumors (10–14). Large-scale studies from The Cancer Genome Atlas (TCGA) and the Molecular Taxonomy of Breast Cancer International Consortium (METABRIC) have reported genomic and transcriptomic analyses on resected IDC and ILC tumors (15,16). The distinguishing genomic feature of ILC is the loss of E-cadherin, a protein that mediates epithelial-specific cell-cell adhesion (17). The loss of E-cadherin in *CDH1* mutants is associated with phosphatidylinositol 3 kinase (PI3K)/Akt pathway activation and epidermal growth factor receptor (EGFR) overexpression, which are major drivers in breast cancer (17). E-cadherin knock-outs of IDC cell lines result in remodeling of transcriptomic membranous systems, greater resemblance to ILCs, and increased sensitivity to IFN-γ-mediated growth inhibition via activation of IRF1 (18).

The National Cancer Institute (NCI) Genomic Data Analysis Network (GDAN) generated assay for transposase-accessible chromatin with high-throughput sequencing (ATAC-seq) data for a subset of TCGA samples (404 patients) (19), including ER+ ILC and IDC tumors. ATAC-seq is a transformative technology for mapping the chromatin-accessible loci genome-wide and identifying nucleosome-free positions in regulatory regions. It needs only ~50,000 cells and is simpler than previous methods, such as DNase-seq (20). Epigenomic changes at the level of chromatin accessibility, potentially linked to distinct differentiation states, might reveal epigenetic reprogramming and developmental origin differences between ER+ ILC and IDC. However, chromatin accessibility landscape differences between ER+ ILC and IDC tumors based on patient samples have not been systematically characterized. Complementing genomic and transcriptomic studies, we mapped the epigenetic heterogeneity in ER+ ILC and IDC with a systematic analysis of chromatin accessibility patterns based on the primary tumor breast cancer TCGA ATAC-seq dataset (21). We defined the compendium of ~190,000 genome-wide cis-regulatory regions in breast cancer ER+ ILC and IDC with 11,762 differentially accessible (DA) peaks between ILC and IDC, which represented 5.98% of total ATAC-seq peaks. EGR1, RUNX3, TP63, STAT6, SOX family, and TEAD family transcription factor (TF) activities were significantly higher in ILC, consistent with their role in the regulation of the extracellular matrix and growth factor signaling pathways, whereas ATF4, PBX3, SPDEF, PITX family, and FOX family TF activities were significantly higher in IDC. The inferred TF activities and context-specific target genes based on ATAC-seq data identified biological pathways that are likely distinct in ER+ ILC vs. IDC. Together, these results provide new insights into ER+ ILC and IDC biology.

## Results

### Chromatin accessibility differences between ER+/HER2- ILC and IDC breast cancer

Epigenetic differences at the level of chromatin accessibility, potentially linked to different differentiation states, could distinguish ER+ ILC and IDC tumors. We characterized tumor cell-intrinsic chromatin accessibility patterns using primary ER+ breast cancer ATAC-seq data from TCGA (19). Using an atlas of 196,546 peaks across all ILC and IDC breast tumors (n=67) (19), we grouped tumors according to their histological subtypes and hormone receptor status: ER+/HER2- ILCs (n=13), ER+/HER2- IDCs (n=30), and ER+/HER2+ IDCs (n=7). Principal component analysis (PCA) of peak read counts showed that all these ER+ tumors were clustered closely, the ER+/HER2- or HER2+ IDCs more closely associated, and the ILCs slightly separated **(Fig. 1A)**. In addition, we found three ER+/HER2- IDC samples and one ER+/HER2+ IDC sample that were outliers. We observed similar patterns and the same outliers through PCA analysis of RNA-seq and reverse phase protein array (RPPA) data (**Supplementary Figure 1A–B**). Because there were few ER+/HER2+ samples, we used ER+/HER2- ILCs (n=13) and ER+/HER2- IDCs (n□=□27) for downstream analyses. Hereafter, we simply denote ILCs vs IDCs omitting ER+/HER2−.

**Figure 1.**
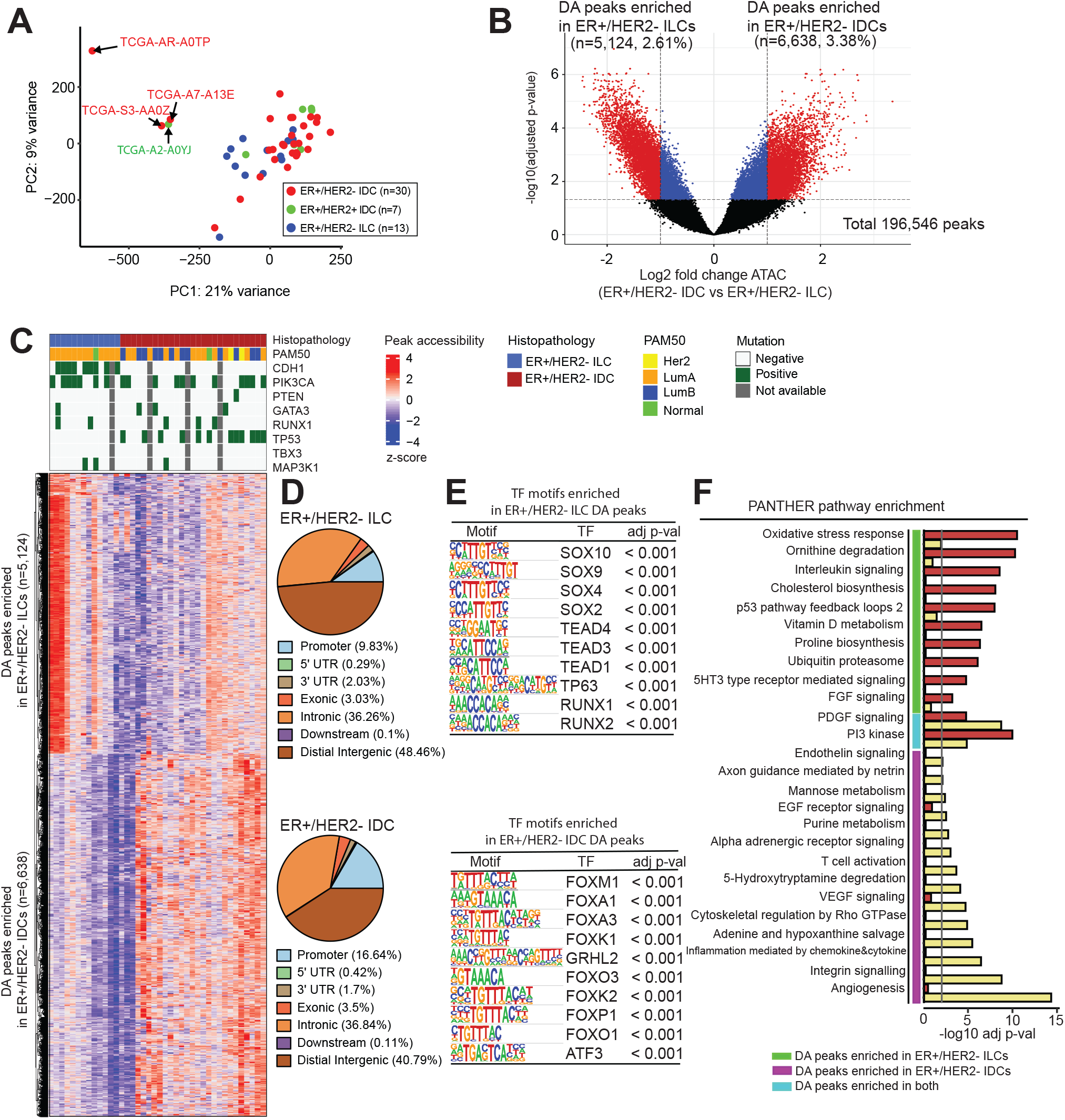
Differential chromatin accessibility between ER+/HER2- ILC and ER+/HER2- IDC. **(A)** PCA of ATAC-seq signal in all peaks (n=196,546). All tumors of ER+ were clustered, but ILC and IDC tumors were slightly separated. The three outliers of ER+/HER2- IDCs and one outlier of ER+/HER2+ IDCs are annotated in the plot. **(B)** Volcano plot of ATAC-seq peaks comparing ILCs (n=13) to IDCs (n=27). Significant peaks with differential chromatin accessibility are highlighted in red. The vertical dotted line indicates an absolute log2 fold change of 1.0 and the horizontal dotted line indicates an FDR-corrected p-value 0.05 criterion; the DA peaks enriched in ILCs (n=5,124) vs IDCs (n=6,638). FDR-corrected p-values were obtained using DESeq2. **(C)** Hierarchical clustering of the 11,762 DA peaks. The significant DA peaks identified in Fig. 1B were aggregated to 11,762 peaks and represented as chromatin accessibility patterns in ILCs and IDCs. Colors represent log2-transformed peak count data and the z-score row was normalized. **(D)** Pie charts show the percentage of DA ATAC-seq peaks (FDR□<□0.05) at the promoter, intronic, intergenic, and exonic regions for ILCs vs. IDCs. **(E)** Enrichment of TF-binding motifs for the subclusters of DA regions of ILCs and IDCs. The top 10 enriched motifs are shown. **(F)** Enrichment of PANTHER pathways for subclusters of DA regions of ILCs and IDCs. In the bar plot, the grey line indicates the significance of the PANTHER pathways (hypergeometric test, adjusted p-value < 0.05). GREAT tool was used to identify the PANTHER pathways overrepresented in the DA peaks.

To understand epigenomic landscape differences between ILCs and IDCs We analyzed differential chromatin accessibility. We found 11,762 DA peaks (absolute log_2_ fold change > 1.0 and adjusted p-value □<□ 0.05) between ILCs and IDCs, which represented 5.98% of all ATAC-seq peaks (**Fig. 1B–C**). Among these, 5,124 peaks (2.61%) showed increased accessibility in ILCs, and 6,638 peaks (3.38%) showed increased accessibility in IDCs. Most of the DA ATAC-seq peaks were at distal intergenic regions (48.46% for ILCs and 40.79% for IDCs); 36.26% for ILCs and 36.84% for IDCs were at introns; 9.83% for ILCs and 16.64% for IDCs were at promoters; and 3.03% for ILCs and 3.5% for IDCs were at exons (**Fig. 1D**).

We used HOMER motif analysis of DA ATAC-seq peaks to identify key TFs driving the expression differences between IDCs and ILCs (22). DA sites in ILCs were highly enriched with binding motifs for the Sry-related HMG box (SOX), TEA Domain (TEAD), runt-related transcription factor (RUNX) family, and TP63 TFs. In contrast, SPDEF and forkhead box (FOX) family binding motifs (e.g., FOXM1, FOXA1, FOXK1) were enriched in IDC sites (**Fig. 1E**). Interestingly, SOX family TFs were major predicted motifs in DA promoter peaks enriched in ILCs, but not in DA distal intergenic peaks (**Supplementary Figure 2A–B**). FOX family TFs were dominant motifs in the DA distal intergenic peaks enriched in IDCs, but not in the DA promoter peaks (**Supplementary Figure 2C–D**). Although SOX-mediated transcription regulation is active in breast cancer (23,24), no association with histological subtypes was reported. Upregulated SOX2 proteins induce chemoresistance in breast cancer cells and promote their stemness property through the recruitment of regulatory T cells (Tregs) to the tumor microenvironment (25,26). SOX4 is an oncogene that promotes PI3K/Akt signaling, angiogenesis, and resistance to cancer therapies in breast tumors; thus, SOX4 is a biomarker for PI3K-targeted therapy (27,28). TEADs interact with transcription coactivator Yes-associated protein (YAP)/transcriptional coactivator with PDZ-binding motif (TAZ), thereby affecting the Hippo pathway that plays a key role in cell proliferation, invasion, and resistance to breast cancer treatment (29,30). TP63, a member of the TP53 gene family, is highly expressed in metaplastic breast cancer (31). SPDEF function depends on the breast cancer subtype (32). SPDEF is a tumor suppressor in triple-negative breast cancer (TNBC) inhibiting tumor invasion and decreasing epithelial-mesenchymal transformation (EMT) (33). In luminal or HER2+ breast cancer, SPDEF is an oncogene (34). FOXA1 proteins enhance hormone-driven ER activity and binding to intergenic regions of DNA in ER+ breast cancer (35). FOXA1 also inhibits EMT and cell growth by modulating E-cadherin, leading to a better prognosis (36).

To identify key biological processes that drive oncogenic gene expression differences between IDCs and ILCs, we analyzed the pathways for DA ATAC-seq peaks using the Genomic Regions Enrichment of Annotations Tool (GREAT) (37). The DA peaks enriched in ILCs and IDCs were associated with different pathways. Oxidative stress response, interleukin signaling, and p53 pathways were overrepresented in ILCs, whereas endothelin signaling, EGF receptor signaling, T cell activation, inflammation, and angiogenesis pathways were overrepresented in IDCs (p-value < 0.01) (**Fig. 1F**). Interestingly, the PI3K pathway was enriched in both ILCs and IDCs. Thus, the epigenomic differences identified distinct TF motif enrichment and biological signatures between ILCs and IDCs.

To correlate alterations in chromatin accessibility with transcriptional output, we integrated ATAC-seq data with RNA-seq data. Consistent with the correlation of global differential accessibility and expression, differential accessibility of individual genes was often associated with significant differential expression (**Fig. 2A–B**). Genes with the greatest differential accessibility between ILCs and IDCs at their promoter, intronic, and nearby intergenic peaks are shown in **Fig. 2B**. For example, FAM83A and ERICH5 were significantly more accessible in IDCs, while FAM189A and SSPN were significantly more accessible in ILCs (**Fig. 2C**). FAM83A is involved in the chemoresistance and stemness of breast cancer through its interaction with the EGFR/PI3K/AKT signaling pathway (38,39). FAM189A is down-regulated in breast cancer (40,41) and SSPN is down-regulated in TNBC (42). Overall, we identified context-specific features, including accessibility and expression patterns associated with IDCs vs. ILCs.

**Figure 2.**
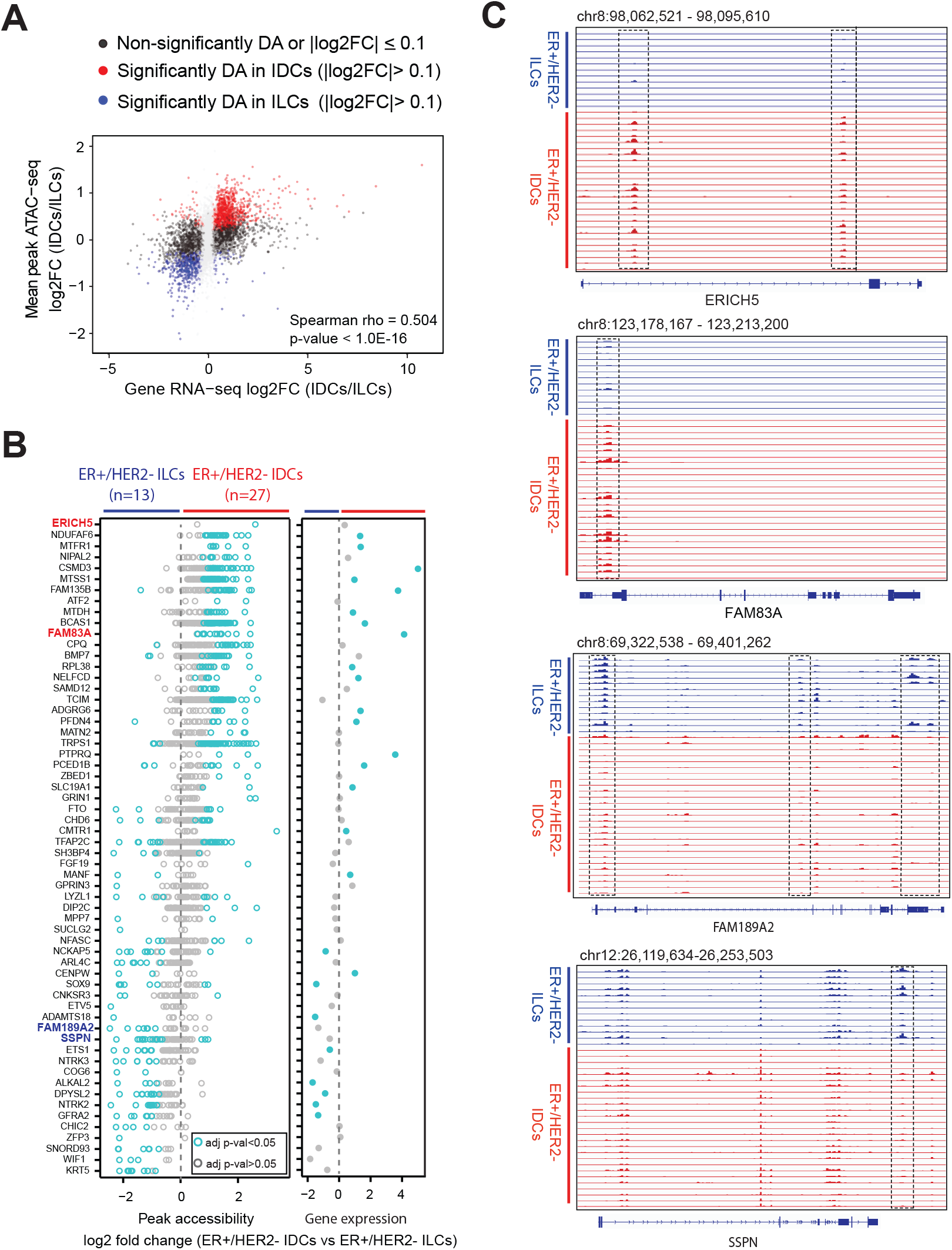
ILC and IDC tumors share a common chromatin state space. **(A)** Scatter plot of differential expression (RNA-seq log2FC, x-axis) and differential accessibility (mean ATAC-seq log2FC over all peaks associated with a gene, y-axis) between IDCs and ILCs tumors. Significantly DA genes are highlighted with red or blue color. **(B)** Differential accessibility and differential expression between ILCs and IDCs. Left: ATAC-seq signal log2 fold change for peaks of significantly DA genes; right: log2 fold change of RNA-seq gene expression (color for significantly decreased/increased individual peaks or genes; adjusted p-value < 0.05). **(C)** The upper panel depicts genome browser tracks (GRCh38) showing chromatin accessibility at ERICH5 and FAM18A2 gene loci in ILCs and IDCs. The lower panel of genome browser tracks shows chromatin accessibility at FAM189A2 and SSPN gene loci which have DA peaks enriched in ILCs. The dotted line boxes highlight the ATAC-seq peaks of DA between ILCs and IDCs. All the track lines have the same y-axis limits and the peak height is scaled over all samples.

### The coordinated activity of many TFs characterizes ILC and IDC tumors

We inferred sample-specific TF motif activities based on genome-wide chromatin accessibility data using CREMA (Cis-Regulatory Element Motif Activities, see Methods). This allowed us to map chromatin accessibility profiles to a lower-dimensional inferred TF activity space, largely preserving the relationships between samples. Inferred activities of 29 TF motifs were significantly associated with histological subtypes by false discovery rate (FDR)-corrected p-value < 0.05 and absolute mean activity difference > 0.035 (**Fig. 3A and Supplement Figure 3**). We found that Early Growth Response 1 (EGR1) (43), TEAD family (TEAD1, TEAD3, and TEAD4), SOX family, (SOX2, SOX4, and SOX8), and RUNX3_BLC11A TFs had significantly higher activities in ILCs than IDCs (**Fig. 3B**). Similarly, FOX family (FOXA1, FOXA3, FOXC2, FOXL1, FOXK1, FOXP2, FOXP3, FOXD3, FOXI1, and FOXF1), Paired Like Homeodomain (PITX family) (PITX1 and PITX2), PBX3, and HSF4 had significantly higher activities in IDCs than ILCs (**Fig. 3C**). EGR1 mRNA is upregulated in ILCs (43), but other TFs have not been studied in the context of ILCs and IDCs. TF activities from the same families were also correlated across samples (**Fig. 3D**). Overall, these results were consistent with the motif enrichment analysis based on the DA peaks in ILCs vs. IDCs (**Fig. 1E**).

**Figure 3.**
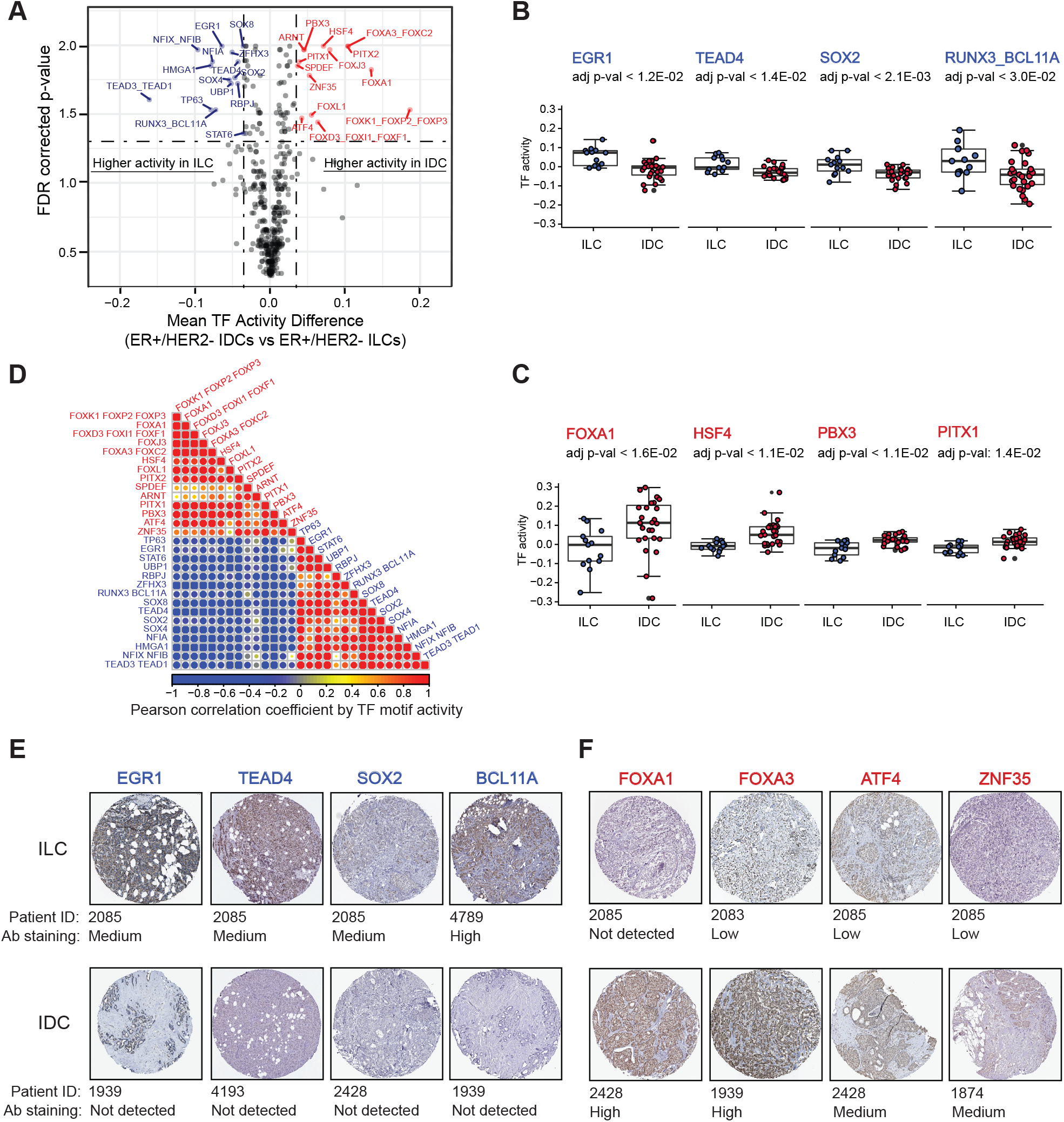
ATAC-seq analysis identifies key TFs in ILC and IDC tumors. **(A)** Inferred TF motif activity differences between ILCs and IDCs. The x-axis is the mean TF activity differences and the y-axis is –log10 (FDR-corrected p-values). Multiple hypothesis testing correction was done using the Benjamini–Hochberg procedure. The vertical dotted line indicates an absolute mean TF activity difference of 0.035 and the horizontal dotted line indicates the FDR-corrected p-value = 0.05 for significant TFs.**(B)** EGR1, TEAD4, SOX2, RUNX3_BCL11A had inferred high TF activities in ILCs. **(C)** FOXA1, FOXA3, ATF4, and ZNF35 had inferred high TF activity in IDCs. The significance of the TF motif activity difference was determined by the Wilcoxon rank-sum test adjusted p-value. **(D)** The Pearson correlation for TF activities in all ILC and IDC tumors. **(E)** Immunohistochemical staining for TF protein expression. EGR1, BCL11A, TEAD4, and SOX2 had high activities in ILCs. Their protein expressions were high or medium in ILCs, but not detected in IDCs. **(F)** FOXA1, FOXA3, ATF4, and ZNF35 had high activities in IDCs. Their protein expression was high or medium in IDCs, but not detected or low in ILCs. All tumors had high or medium ER expression, but HER2 expression was not detected or low.

To determine whether TF activities were associated with protein expression, we compared immunohistochemically (IHC) stained images for available TFs between ILCs and IDCs obtained from the Human Protein Atlas (HPA) database (44). The HPA tissue images of breast tumors provide histological subtype but not hormone receptor subtype information. Therefore, we used only ILC or IDC images that had ER high/medium staining intensity and HER2 low/not detected staining intensity (**Supplementary Figure 4**). The IHC images demonstrated that EGR1, TEAD4, SOX2, and BCL11A proteins were highly expressed in ILCs, but were not detected in IDCs (**Fig. 3E**) consistent with the increased TF activities in ILCs. Likewise, the IHC images of FOXA1, FOXA3, ATF4, and ZNF35 showed medium or high protein expression in IDCs but were not detected or showed low expression in ILCs (**Fig. 3F**). Overall, the staining images showed that increased TF activities in ILCs or IDCs were associated with protein expression in the corresponding histological subtypes.

We looked for functional evidence of IDC- and ILC-specific TF regulators using published breast cancer genome-wide ‘‘dropout” screens using pooled small hairpin RNA (shRNA) libraries (45). The dataset included three ER+ ILC cell lines and 11 ER+ IDC cell lines (HER2 type data was not available). We ran **s**mall **i**nterfering RNA (siRNA)/shRNA **m**ixed-**e**ffect **m**odel (siMEM) for three ER+ ILC cell lines vs. other breast cancer cell lines or for 11 ER+ IDC cell lines vs others to calculate context-specific essentiality scores for IDC- or ILC-specific TFs. **Supplementary Table 1** lists the essentiality scores of the TFs in ER+ ILC (15/18 TFs) or ER+ IDC (15/19 TFs) cell lines. We identified RUNX3, SOX4, TEAD3, UBP1, NFIA, and BCL11A as essential for ER+ ILC cell proliferation, and FOXA1 and SPDEF as essential for ER+ IDC cell proliferation (FDR < 0.2). RUNX3 inhibits estrogen-dependent proliferation by targeting ERα in breast cancer cell lines and functions as a tumor suppressor, but its role has not been defined (46). Interestingly, we identified FOXA1 and SPDEF as the top essential TFs for ER+ IDC cells. The original genome-wide shRNA screening also identified FOXA1 and SPEDEF as the top luminal/HER2 essential genes out of 975 essential genes (45). In ER+ breast cancer cell lines, FOXA1 inhibits cell growth by inducing E-cadherin expression and suppressing ER pathway activity, which suggests that FOXA1 can be a favorable prognostic marker in human breast cancer (36,47,48). SPDEF expression is also enriched in luminal tumors and promotes luminal differentiation and survival of ER+ cells (49).

### Gene sets for IDC- and ILC-specific TFs display coherent functions and are consistent with gene expression changes

**Tables 1** and **2** summarize the enriched canonical pathways for the target genes of the TFs associated with ILCs or IDCs. Interestingly, most TFs with high activities in ILCs, including EGR1, HMGA1, NFIX_NIFB, RBPJ, SOX family, TEAD family, TP63, UBP1, and ZFHX3, were associated with genes encoding extracellular matrix (ECM)-associated proteins for structure or remodeling. IDC-specific TFs including ARNT, ATF4, and ZNF35 were associated with PI3K or IL2 signaling mediated by PI3K. We next examined TF target gene expression in IDCs and ILCs using TCGA and METABRIC gene expression data. **Fig. 4** shows the cumulative distribution of expression changes between ILC- or IDC-specific TF activities for predicted targets based on ATAC-seq data. The TF regulators identified for ILC and IDC were associated with the upregulation of their targets. There was significant upregulation of motif-based targets of TFs based on ATAC-seq relative to all genes in the TCGA and METABRIC tumor data (p-value < 1e-3, Kolmogorov-Smirnov test). Thus, ILCs and IDCs are associated with different TFs, and the TFs regulate target gene expression and biological pathways specific for ILCs vs. IDCs.

**Table 1:**
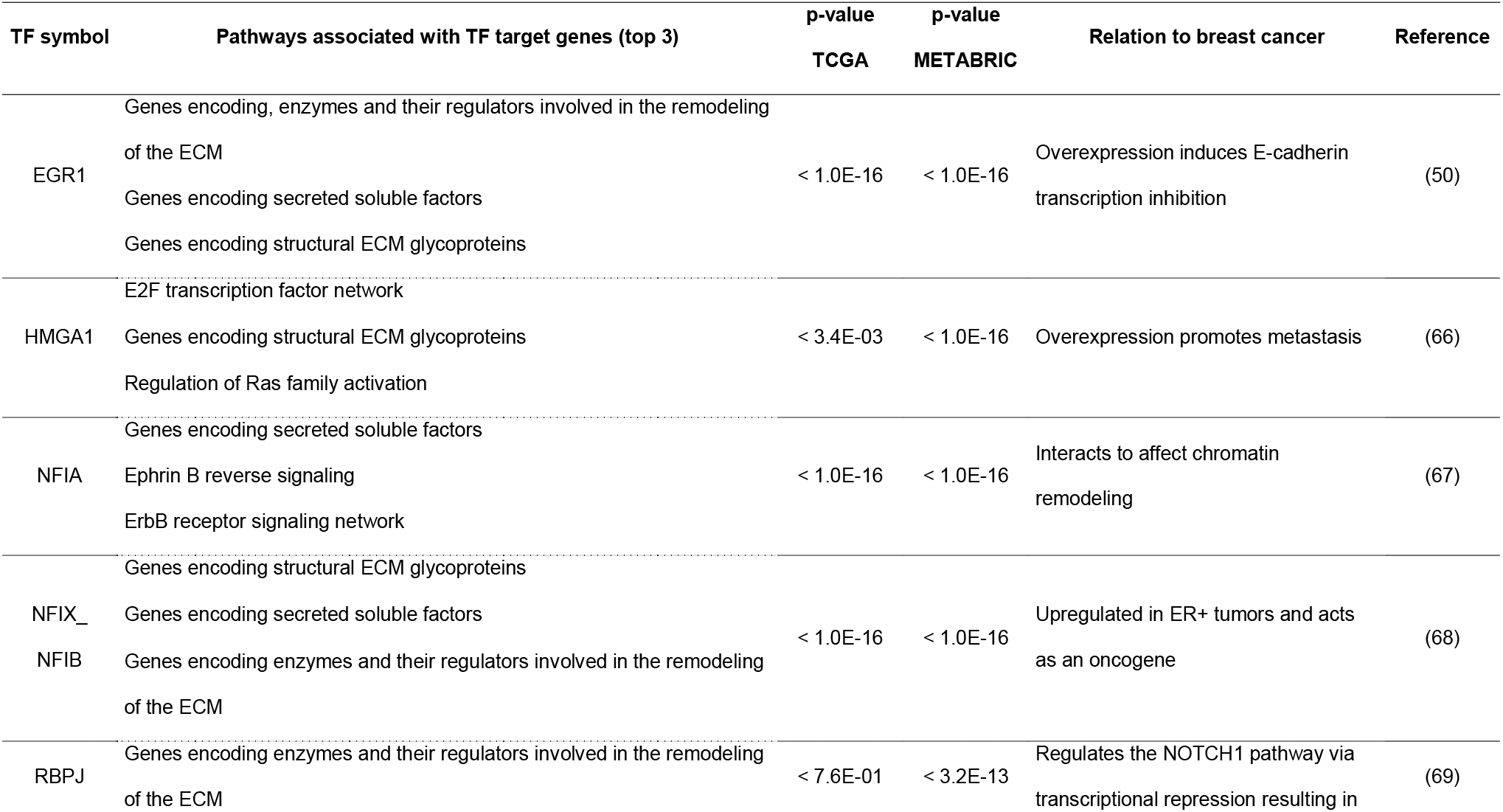
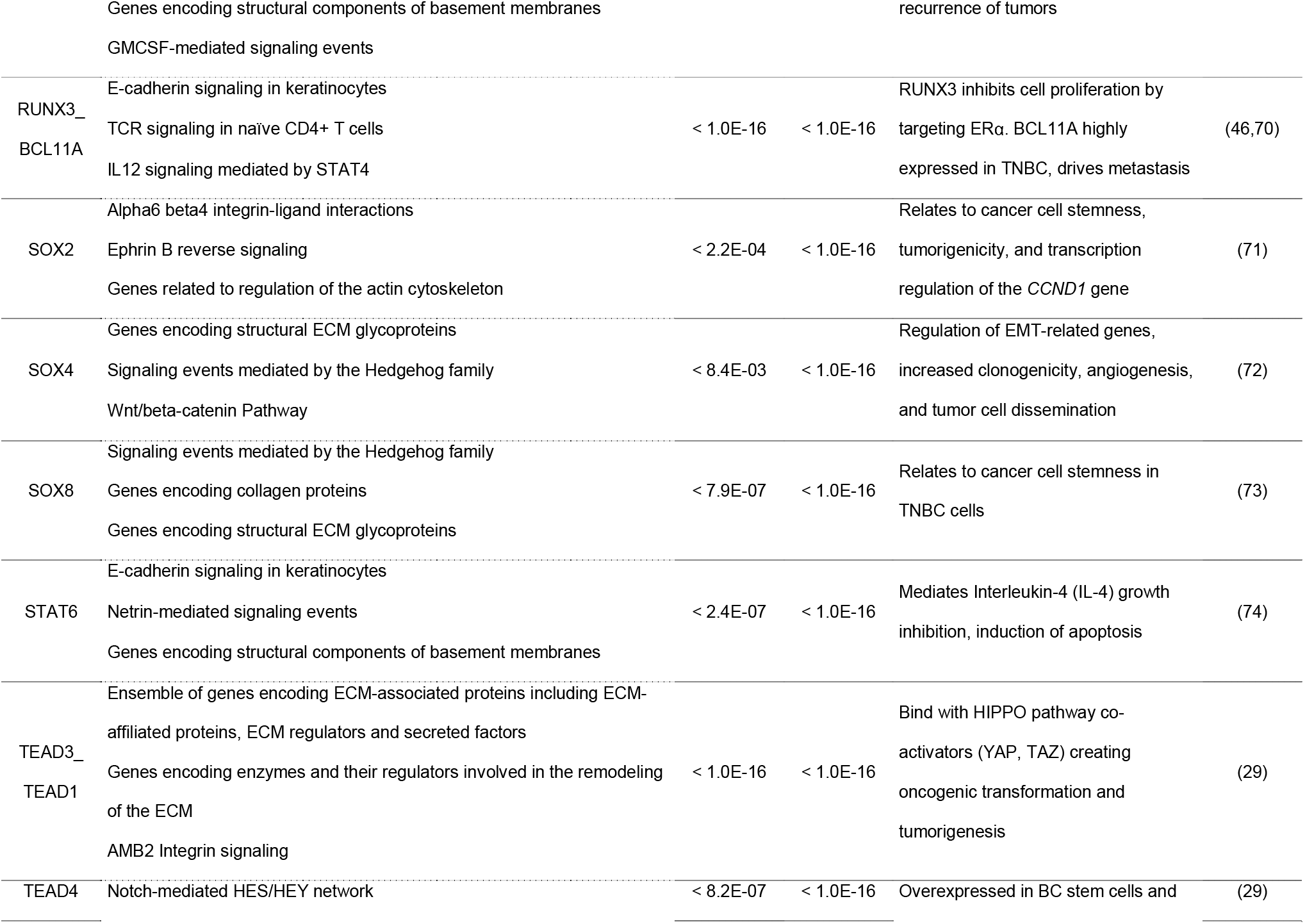
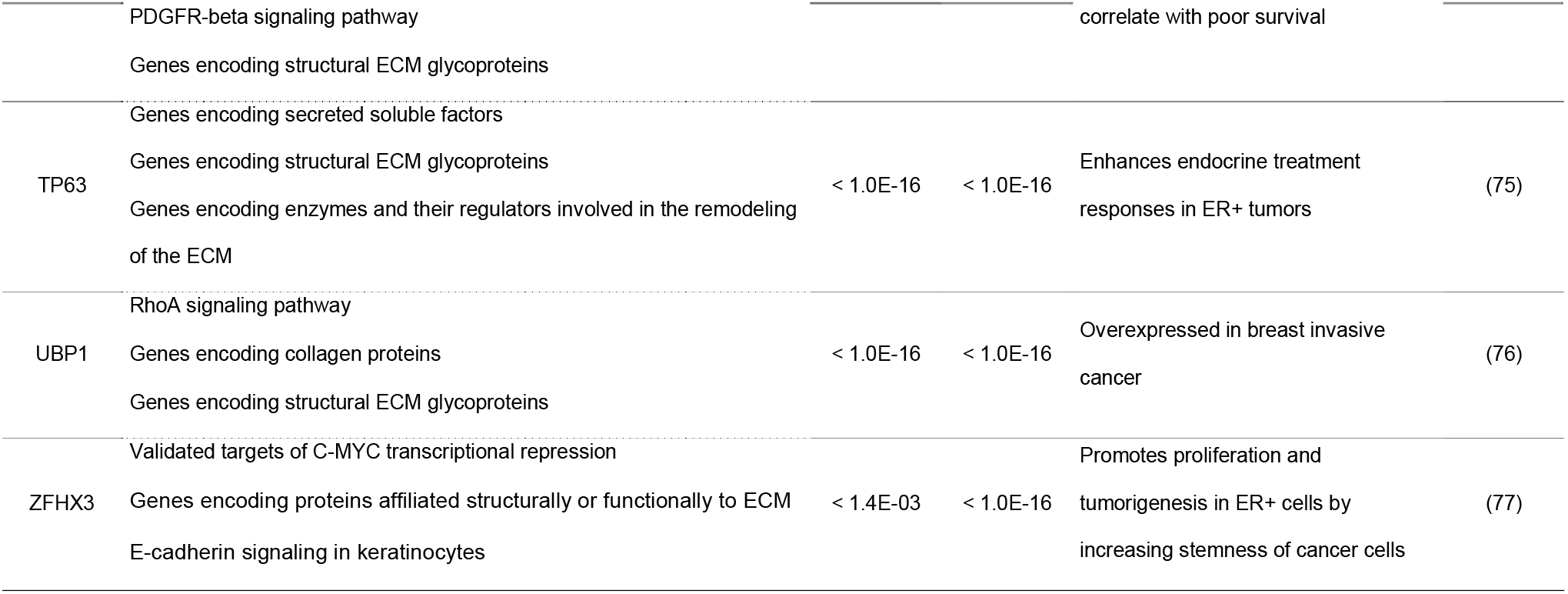
Candidate TF regulators selected at 5% FDR for ILC. Functional annotations were determined from terms overrepresented from the canonical pathway gene sets associated with the candidate regulator. The p-values are from the Kolmogorov-Smirnov (K-S) test between the target and the background distributions for TCGA and METABRIC datasets.

**Table 2:**
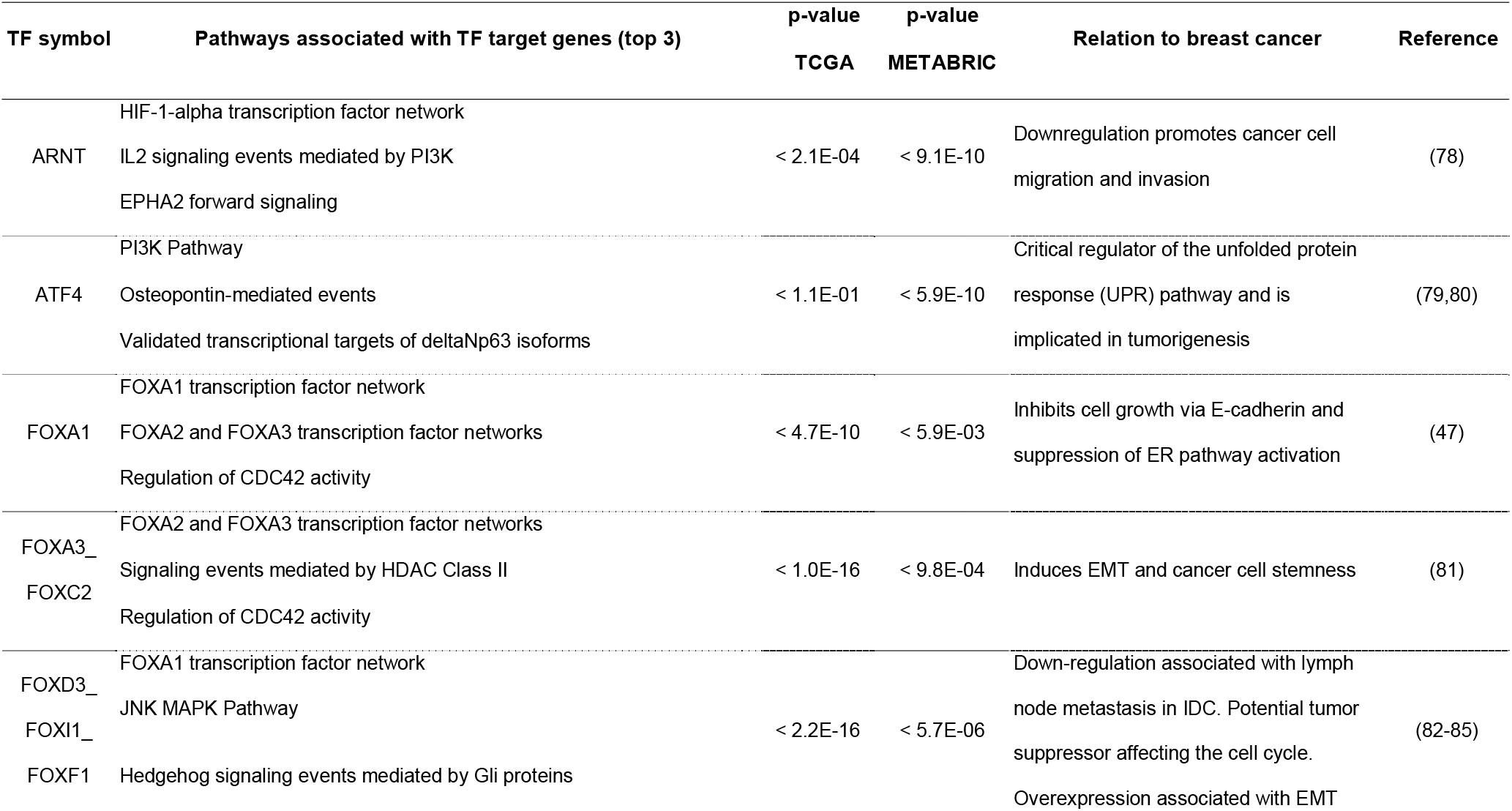
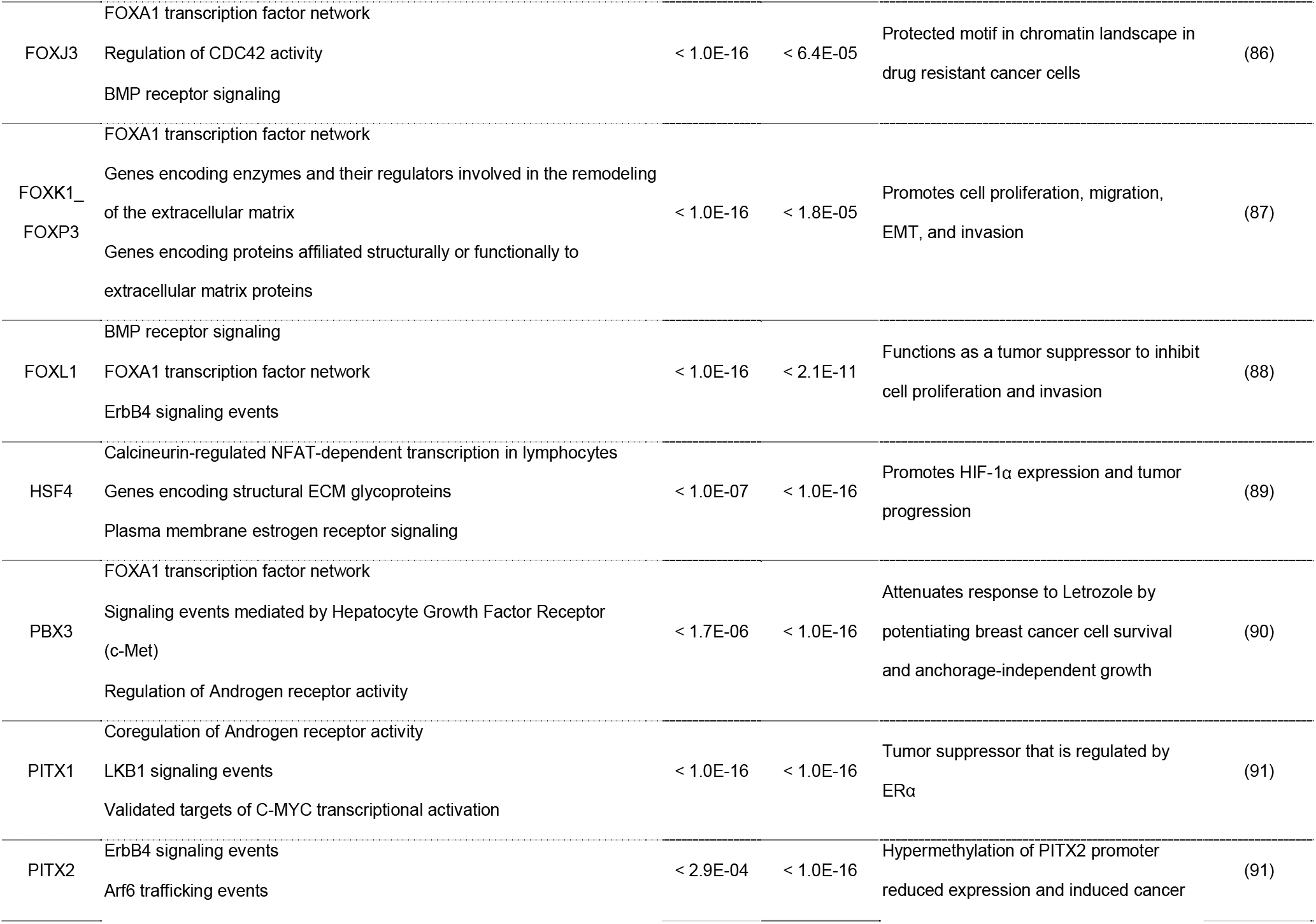
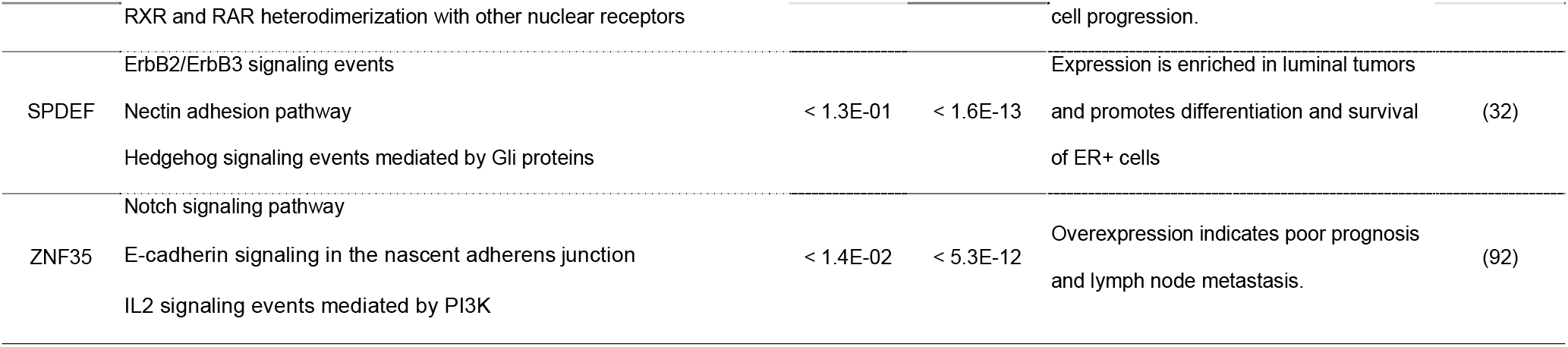
Candidate TF regulators selected at 5% FDR for IDC. Functional annotations were determined from terms overrepresented from the canonical pathway gene sets associated with the candidate regulator. The p-values are from the Kolmogorov-Smirnov (K-S) test between the target and the background distributions for TCGA and METABRIC datasets.

**Figure 4.**
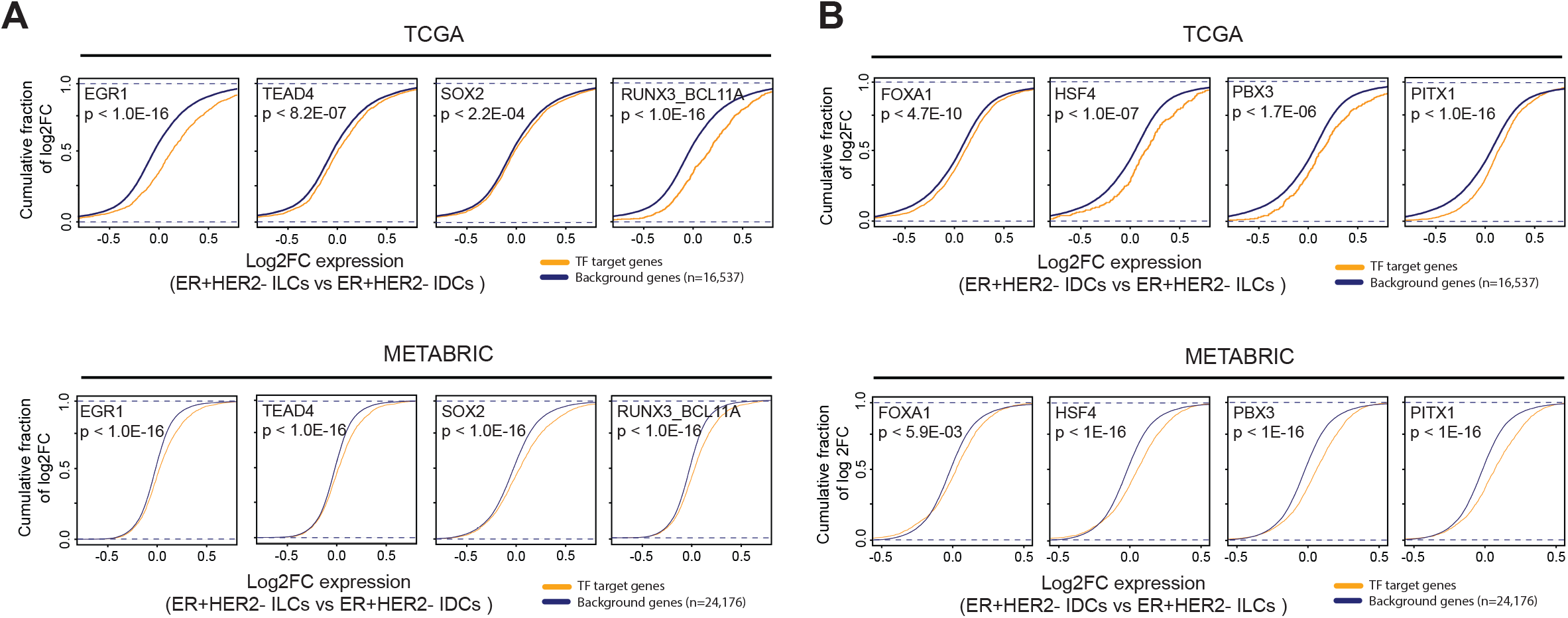
Gene sets for candidate ILC- and IDC-specific TFs display coherent functional annotations and consistent expression changes in tumors. **(A)** Targets of EGR1, TEAD4, SOX2, and RUNX3_BCL11A, ILC-specific candidate TFs, showed significant upregulation in ILC tumors relative to IDC tumors (p-value < 1e-3, Kolmogorov-Smirnov test) compared to background genes. The upper panel depicts the upregulation of TF target genes expression in TCGA RNA-seq data and the bottom panel depicts METABRIC microarray data. **(B)** Targets of FOXA1, HSF4, PBX3, and PITX1, IDC-specific candidate TFs showed significant upregulation of expression in TCGA and METABRIC data. The background genes were all genes identified in the gene expression dataset after removing low or non-expression genes. The yellow lines are empirical Cumulative Distribution Functions (eCDF) for the target gene log2 fold changes between ILCs and IDCs. The blue lines are CDFs for background gene log2 fold changes between ILCs and IDCs. The p-values are from the Kolmogorov-Smirnov (K-S) test between the target and the background distributions.

## Discussion

Many studies have examined the distinct molecular features and prognostic outcomes for ILC vs. IDC tumors. We provide here the first comprehensive genome-wide chromatin accessibility landscape analysis of ER+ ILC and IDC using primary breast cancer TCGA ATAC-seq data. We identified a chromatin accessibility signature, TFs, and biological pathways specific to ER+ ILC and IDC tumors. TFs (e.g. EGR1, SOX, and TEAD family) involved in ECM interactions, developmental pathways had higher activity in ILC compared to IDC. The differences in activities of TFs in ILCs vs. IDCs based on chromatin accessibility were also consistent with TF protein expression and upregulation of TF target gene expression.

The altered TF activities associated with these histological subtypes have a direct relationship to the biological presentation of the resulting tumors and their diagnosis. For example, EGR1 can be activated via the MAPK signaling pathway through stimulation by reactive oxygen species (50) consistent with our pathway enrichment analysis in ILCs (**Fig. 1F** and **Table 1**). Further, EGR1 contributes to tumor invasion and metastasis in ovarian cancer cells by activating the expression of SNAIL and SLUG which are E-cadherin transcriptional inhibitors (51). The TEAD family, specifically TEAD4, has been shown to bind with the oncogenic TF KLF5 and in turn induce transcription of fibroblast growth factor binding protein 1 (FGFBP1), which is promoting cell proliferation through expansion of the fibroblast cell type in TNBC (52). Lastly, the SOX family TFs are critical regulators of developmental processes and contribute to tumor development and progression. SOX TFs have been shown to act in both an oncogenic capacity and as a tumor suppressor (24). SOX2 and SOX9 are shown to interact during increased cancer stem cell content and the development of drug resistance. SOX2 increase in association with estrogen reduction reduces the expression of the SOX9. SOX9 is known to work downstream of SOX2 to control the luminal progenitor cell content resulting in increased tumor initiation, drug resistance, and poor prognosis (53). Our results suggest that the potential role of these TFs in ILC and IDC merits further investigation.

## Conclusions

This study provides the first in-depth characterization of the genome-wide chromatin accessibility landscape of ER+ ILC and IDC primary tumors samples. We identified several differences in the epigenomic profiles between ILC and IDC and highlighted potentially clinically relevant pathways. Our deep analyses of ATAC-seq data generated a global regulatory network with the corresponding TFs in IDC and ILCs that could provide useful clinical insights into the differences between these two histological subtypes.

## Methods

### Data and preprocessing

#### TCGA breast cancer data

We downloaded TCGA breast cancer (BRCA) ATAC-seq raw bam files (n=75) and RNA-seq raw fastq.gz files from NCI Genomic Data Commons (GDC) data portal (https://portal.gdc.cancer.gov) (54). Breast cancer peak calls and bigwig files of ATAC-seq profiles were downloaded from https://gdc.cancer.gov/about-data/publications/ATACseq-AWG. For the hormone receptor subtypes of TCGA BRCA tumors, we followed the annotation data provided by the TCGA ATAC-seq data publication, Supplementary Data 1 (21). The RNA-seq read count and reverse phase protein array (RPPA, replicate-base normalization) data were downloaded from Xena Functional Genomics Explorer (https://xenabrowser.net/hub/) GDC and TCGA hub, respectively.

#### METABRIC breast cancer data

We downloaded METABRIC microarray data from cBioPortal (https://www.cbioportal.org/study/summary?id=brca_metabric) (55).

#### Human Protein Atlas

The Human Protein Atlas (https://www.proteinatlas.org) is a public resource that extracts and analyzes information, including images of immunohistochemistry (IHC), protein profiling, and pathologic information, from specimens and clinical material from cancer patients to determine global protein expression (56). Here, we compared the protein expression of available TFs in ILC and IDC tissues by IHC image.

#### Breast cancer cell lines shRNA screen

To identify and validate the TFs essential in ER+ ILC and ER+ IDC cell lines, we accessed the data for whole-genome small hairpin RNA (shRNA) ‘‘dropout screens” on three ER+ ILC and 11 ER+ IDC breast cancer cell lines (45) (GSE74702).

### Differential peak accessibility

Reads aligning to atlas peak regions (hg19) were counted using the countOverlaps function of the R packages, GenomicAlignments (v1.30) (57) and GenomicRanges (v1.46.1) (57). To remove the bias created by low count peaks, we filtered 19,364 peaks with mean counts of less than 30 across all samples. Differential accessibility of peaks was calculated using DESeq2 (v1.34.0) (58). DA peaks were defined as significant if they had an adjusted p-value < 0.05 and the magnitude of the DESeq-normalized counts changed by a stringent factor of one or more between ER+HER2- ILC and ER+/HER2- IDC. The significant DA peaks were aggregated and represented in the hierarchical clustering heatmap using the DESeq size-factor normalized read counts and the ‘complete’ distance metric for the clustering algorithm. We used ChiPseeker (59) and clusterProfiler (60) R packages for peak region annotation and visualization of peak coverage over chromosomes.

### ATAC-seq peak clustering

For visualization of ER+/HER2- ILC, ER+/HER2- IDC, and ER+/HER2+ IDC tumors by PCA, we used DESeq2 (v1.34.0) (58) to fit multi-factorial models to ATAC-seq read counts in peaks. We used all peak counts and generated DESeq2 models including factors for hormone receptor subtypes (ER +/- and HER2 +/-) and histological classes (ILC vs IDC). Then, we calculated a variance stabilizing transformation from the DESeq2 model and performed PCA.

### De novo TF motif analysis

The HOMER v.4.11.1 utility findMotifsGenome.pl (22) was used to identify the top 10 TF motifs enriched in differential accessible peaks. We set 100-bp-wide regions around the DA peak summits with hg19 as the reference genome. We generated custom background regions with a 150-bp-wide range around the peak summits. The top motifs were reported and compared to the HOMER database of known motifs and then manually curated to restrict them to TFs that are expressed based on RNA-seq data and to similar motifs from TFs belonging to the same family.

### Pathway enrichment analysis

We used GREAT (Genomic Regions Enrichment of Annotations Tool, v1.26) to associate the subcluster of the DA peaks to genes and used pathway analysis to identify the functional significance of the DA peaks (37).

### TF essentiality analyses in ER+ ILC and ER+ IDC cell lines

We used **s**mall **i**nterfering RNA (siRNA)/shRNA **m**ixed-**e**ffect **m**odel (siMEM) (45) to score the screening results of the TFs and identify their significant context-specific essentiality between ER+ ILC and ER+ IDC from the shRNA screening data. The significantly essential TFs had an FDR q-value < 0.2 in the siMEM results. The annotation data for ER subtype and histological types in the breast cancer cell lines are available at https://github.com/neellab/simem.

### Cis-Regulatory Element Motif Activity analysis

We used the CREMA (Cis-Regulatory Element Motif Activities, https://crema.unibas.ch/) to analyze genome-wide DNA-accessibility, calculate TF motif activities, and identify active cis-regulatory elements (CREs). CREMA first identifies all CREs genome-wide that are accessible in at least one sample, quantifies the accessibility of each CRE in each sample, predicts TF binding sites (TFBSs) for hundreds of TFs in all CREs, and then models the observed accessibilities across samples in terms of these TFBS, inferring the activity of each TF in each sample. A Wilcoxon rank sum test was used to compare TF activities and assess the association between TF and histologic subtypes. Then, the resulting p-values were adjusted for multiple hypothesis testing (across TFs). This analysis was visualized with a scatterplot where the x-axis represents mean TF activity difference, and the y-axis represents FDR-corrected p-value. The significant TF motifs were selected by absolute mean TF activity difference > 0.035 and FDR-corrected p-value < 0.05.

The TF targets identified by CREMA are CREs, not genes directly. After identifying TF target CREs, the gene-CRE association probabilities are calculated on the basis of distance to transcription start sites (TSSs) of gene within +/- 1,000,000 bp, using a weighing function. The weighing function quantifies the prior probability that a CRE will regulate a TSS at distance d and is a mixture of two Lorentzian distributions with length-scales 150 bp (corresponding to promoter regions) and 50Kb (corresponding to enhancer regions). This weighing function is used to weigh log-likelihood score per possible CRE-TSS interaction. The target gene score is a sum of the log-likelihood scores of all CREs associated with the gene weighted with the association probability. Then, the scores were used to predict over-represented canonical pathways in the TF’s target genes.

### Differential gene expression analysis

We ran DESeq2 on the TCGA RNA-seq read count data between ER+/HER2- ILCs (n=100) and ER+/HER2- IDCs (n=297), which include all available tumors for hormone receptor and histological subtypes. We used the Limma (v3.48) (61) package to calculate the log2 fold change of differentially expressed genes between ER+/HER2- ILCs (n=121) and ER+/HER2- IDCs (n=1,030) for the METABRIC dataset.

We calculated the cumulative distribution of expression changes for the target genes and background genes and ran the Kolmogorov-Smirnov (K-S) statistic to quantify the distance between empirical cumulative distribution function (eCDF) of target genes and cumulative distribution function (CDF) of background genes and determine its significance. We used all 16,537 genes as background genes after removing genes with low mean counts across samples.

### Statistical analysis and data visualization

All statistical analyses were performed using R version 4.1.1 (R Foundation for Statistical Computing, Vienna, Austria) (62). Heatmaps were generated using the R package ComplexHeatmap v2.10.0 (63). Graphs were generated using the R package ggplot2 v3.3.5 (64). Genome track images were generated using the IGV (v2.11.1) (65). P-values in multiple comparisons were adjusted using the Benjamini-Hochberg (BH) method.

## Abbreviations

ATAC-seq: assay for transposase-accessible chromatin with sequencing
CDF: cumulative distribution function
CRE: cis-regulatory elements
CREAM: Cis-Regulatory Element Motif Activities
DA: differentially accessible
eCDF: empirical cumulative distribution function
ECM: extracellular matrix
EGFR: epidermal growth factor receptor
EGR1: Early Growth Response 1
EMT: epithelial-mesenchymal transformation
ERa: estrogen receptors alpha
FDR: false discovery rate
FOX: forkhead box
GDAN: Genomic Data Analysis Network
GREAT: Genomic Regions Enrichment of Annotations Tool
HER2: human epidermal growth factor receptor 2
HPA: Human Protein Atlas
IDC: invasive ductal breast carcinoma
IHC: immunohistochemistry
ILC: invasive lobular breast carcinoma
K-S: Kolmogorov-Smirnov
METABRIC: Molecular Taxonomy of Breast Cancer International Consortium
NCI: National Cancer Institute
PCA: principal component analysis
PI3K: phosphatidylinositol 3 kinase
PR: progesterone receptors
RPPA: RNA-seq and reverse phase protein array
RUNX: runt-related transcription factor
shRNA: small hairpin RNA
siMEM: small interfering RNA (siRNA)/shRNA mixed-effect model
SOX: Sry-related HMG box
TAZ: transcriptional coactivator with PDZ-binding motif
TCGA: The Cancer Genome Atlas
TEAD: TEA Domain
TF: transcription factor
TNBC: triple-negative breast cancer
Tregs: regulatory T cells
TSS: transcription start sites
YAP: yes-associated protein

## Ethics declarations

### Ethics approval and consent to participate

Not applicable.

### Consent for publication

Not applicable.

### Availability of data and materials

This study includes no data or code deposited in external repositories.

### Author contributions

S.L. performed all the computational experiments, analyzed results, and helped to write the manuscript. H.U.O. conceived the project, advised on the analysis, and supervised the research and wrote the manuscript.

## Acknowledgments

The results published here are in whole or part based on the data generated by The Cancer Genome Atlas project established by the NCI and NHGRI (accession number: phs000178.v7p6). Information about TCGA and the investigators and institutions that constitute the TCGA research network can be found at http://cancergenome.nih.gov/. We thank Steffi Oesterreich, Adrian Lee, Jacqueline Bromberg, Jing Hong Wang, Micheal Gatza, Devin Dikec, and Kristi Rothermund for helpful discussions and Mikhail Pachkov and Erik van Nimwegen for their help in CREMA analysis.

