## Supplementary Information for "Chromatin accessibility landscape and active transcription factors in primary human invasive lobular and ductal breast carcinomas"

**Supplementary Figures**

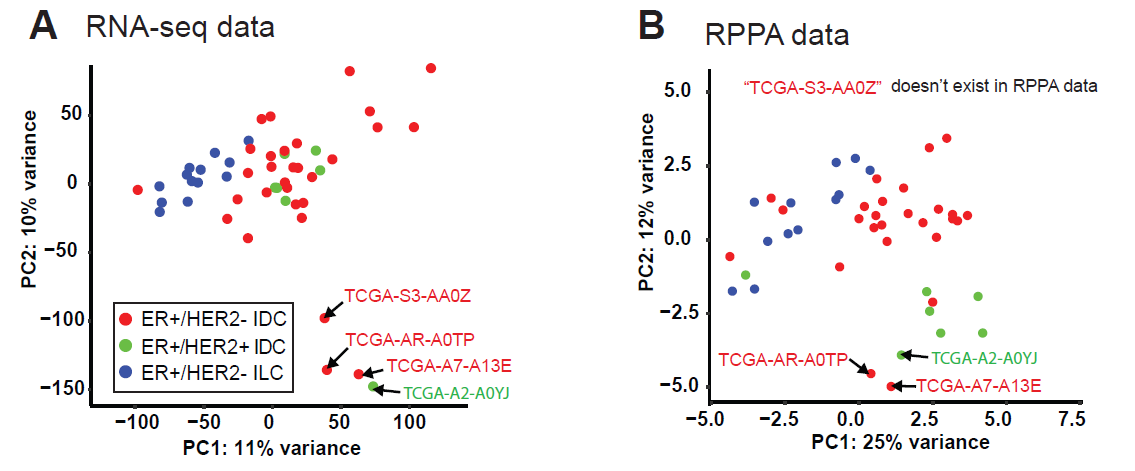

**Supplement Figure 1**. **PCA of gene and protein expression data**. **(A)** ER+ tumor clustering by TCGA RNA-seq data (16,418 genes after removing low read count genes) or **(B)** by Reverse Phase Protein Array (RPPA) data (132 proteins).

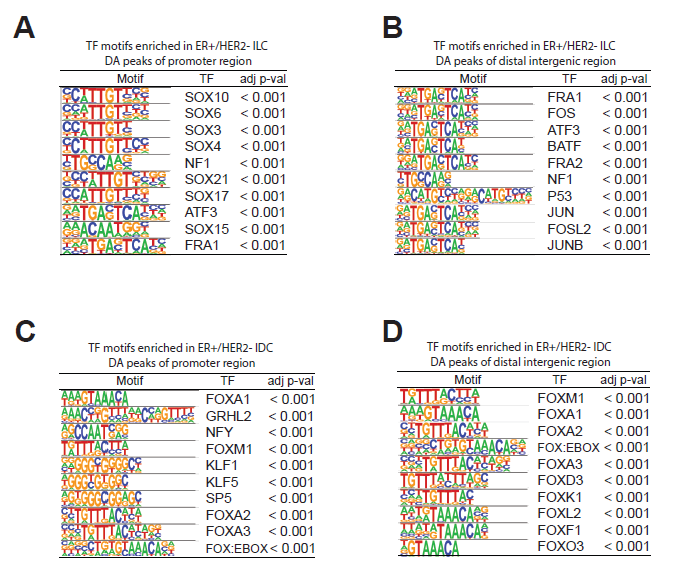

**Supplement Figure 2.** **Enrichment of TF-binding motifs in promoter or distal intergenic regions for ILCs and IDCs.** **(A-B)** Enrichment of TF-binding motifs per the promoter region (491 peaks) or distal intergenic region (2,488 peaks) enriched in ILCs. **(C-D)** Enrichment of TF-binding motifs per the promoter region (1,075 peaks) or distal intergenic region (2,729 peaks) enriched in IDCs. The top 10 most enriched motifs were selected and displayed.

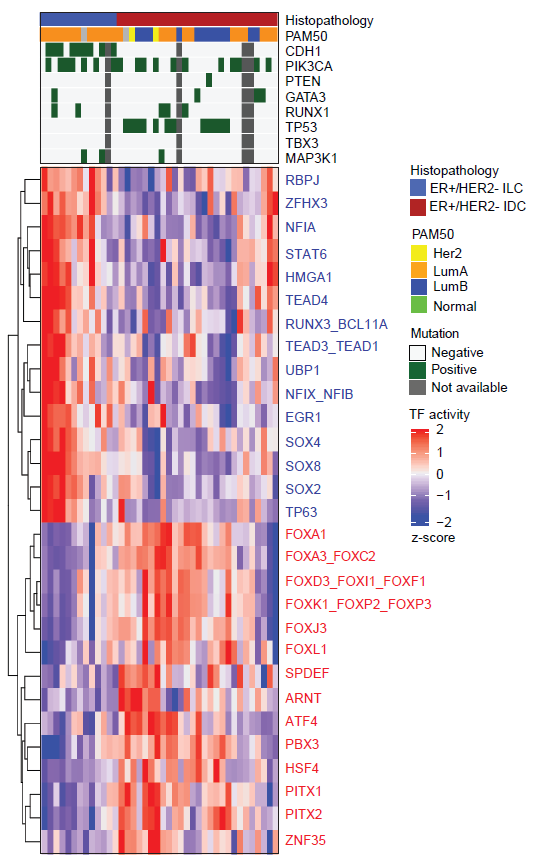

**Supplementary Figure 3.** **Heatmap for TF activities.** TFs significantly associated with ILCs (15 TFs in blue) and IDCs (14 TFs in red) (an absolute mean TF activity difference > 0.035 and the FDR-corrected p-value = 0.05).

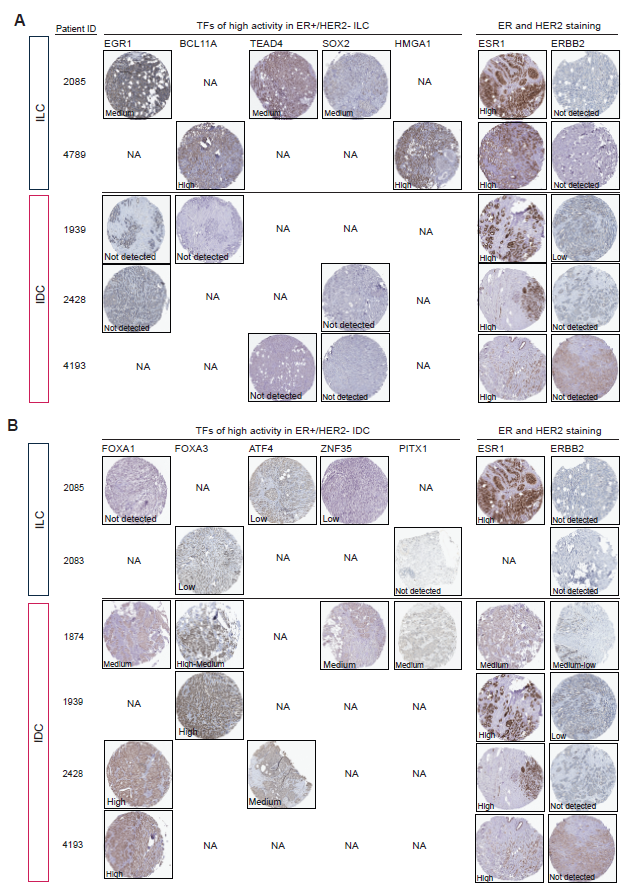

**Supplement Figure 4**. **Immunohistochemical staining for TF protein expression and corresponding ER and HER2 staining. (A)** EGR1, BCL11A, TEAD4, SOX2, and HMGA1, of which high activities in ILCs, have the protein expression high or medium in ILCs, but not detected in IDCs. **(B)** FOXA1, FOX3, ATF4, ZNF35, PITX1, of which high activities in IDCs, have the protein expression high or medium in IDCs, but not detected or low in ILCs. NA, tumor samples of staining images are not available.

**Supplementary Table**

**Supplementary Table 1. List of ER+ ILC or ER+ IDC specific TF genes and essentiality scores derived from siMEM analysis.**

The p-value/FDR derived for each TF according to published screens (GSE73526) using siMEM.

|  | Gene | siMEM score difference | p-value | FDR |
| --- | --- | --- | --- | --- |
| TFs of high activity in ILCs | **RUNX3** | **-0.694** | < 1x10^-16^ | 0.002 |
|  | **SOX4** | **-0.264** | 0.002 | 0.013 |
|  | **TEAD3** | **-0.244** | 0.015 | 0.054 |
|  | **UBP1** | **-0.392** | 0.015 | 0.054 |
|  | **NFIA** | **-0.237** | 0.055 | 0.151 |
|  | **BCL11A** | **-0.173** | 0.065 | 0.151 |
|  | **TEAD1** | 0.239 | 0.094 | 0.189 |
|  | STAT6 | -0.138 | 0.141 | 0.246 |
|  | ZFHX3 | 0.196 | 0.209 | 0.348 |
|  | RBPJ | 0.195 | 0.286 | 0.445 |
|  | NFIB | -0.070 | 0.343 | 0.480 |
|  | NFIX | 0.195 | 0.388 | 0.494 |
|  | HMGA1 | -0.081 | 0.550 | 0.642 |
|  | EGR1 | 0.038 | 0.803 | 0.865 |
|  | TEAD4 | -0.024 | 0.899 | 0.899 |
| TFs of high activity in IDCs | **FOXA1** | **-0.567** | < 1x10^-16^ | < 1x10^-16^ |
|  | **SPDEF** | **-0.480** | 0.001 | 0.008 |
|  | **PBX3** | 0.255 | 0.010 | 0.052 |
|  | **HSF4** | 0.183 | 0.051 | 0.192 |
|  | FOXJ3 | 0.099 | 0.106 | 0.317 |
|  | FOXP3 | 0.098 | 0.137 | 0.343 |
|  | FOXF1 | 0.127 | 0.181 | 0.388 |
|  | FOXC2 | -0.095 | 0.263 | 0.494 |
|  | PITX1 | -0.048 | 0.323 | 0.538 |
|  | PITX2 | 0.037 | 0.538 | 0.808 |
|  | ATF4 | 0.004 | 0.968 | 0.993 |
|  | FOXI1 | -0.018 | 0.815 | 0.993 |
|  | FOXL1 | 0.004 | 0.962 | 0.993 |
|  | ZNF35 | 0.000 | 0.993 | 0.993 |
|  | FOXD3 | 0.027 | 0.778 | 0.993 |

o Yellow highlight: FDR < 0.2

o Blue text: Essential genes in ER+ ILC cell lines

o Red text: Essential genes in ER+ IDC cell lines
